# Increasing number of long-lived ancestors associates with up to a decade of healthspan extension and a healthy metabolomic profile in mid-life

**DOI:** 10.1101/2022.09.08.507098

**Authors:** Niels van den Berg, Mar Rodríguez-Girondo, Ingrid K van Dijk, P. Eline Slagboom, Marian Beekman

**Author notes:** corresponding author, **Corresponding author:** Name: Niels van den Berg, Address: Albinusdreef 2, 2333 ZA Leiden the Netherlands. jointly supervising authors (P. Eline Slagboom, Marian Beekman).

## Abstract

Globally, the lifespan of populations increases but the healthspan is lagging behind. Previous research showed that survival into extreme ages (longevity) clusters in families as illustrated by the increasing lifespan of study participants with each additional long-lived family member. Here we investigate whether the healthspan in such families follows a similar quantitative pattern using three-generational data from two databases, LLS (Netherlands), and SEDD (Sweden). We study healthspan in 2,143 families containing index persons and two ancestral generations, comprising 17,539 persons with 25 follow-up years. Our results provide strong evidence that an increasing number of long-lived ancestors associates with up to a decade of healthspan extension. Further evidence indicates that members of long-lived families have a delayed onset of medication use, multimorbidity and, in mid-life, healthier metabolomic profiles than their partners. We conclude that in longevity families, both lifespan and healthspan are quantitatively linked to ancestral longevity, making such families highly suitable to identify protective mechanisms of multimorbidity.

## Main

The human life expectancy has doubled over the past two centuries^1^, reaching 82.1 years in Western European countries^2^. Although people started to live longer, the time spent in good physical and cognitive health did not rise similarly^2^. In fact, over 70% of the 65 year olds have at least one disease and over 50% have multimorbidity (2 disease or more)^3^. In contrast to the general population, some persons seem to become exceptionally old with a significantly lower chronic age-related disease burden (e.g. high blood pressure, malignancies, and type 2 diabetes) than the general population^4–17^. Moreover, the children of these exceptionally old persons have a delayed first disease onset^11,14,18,19^. These observations are mostly based on cross-sectional designs^4,10–14,17–20^, so prospective studies into the development of first diseases and (multi)morbidity are needed. The study of long-lived families is important as they likely harbor gene-environment interactions which beneficially regulate molecular pathways involved in longevity, resistance to disease, resilience to negative side-effects of treatment and therefore healthy aging^8,21^.

In our previous work we used data from millions of subjects in contemporary medical and historical family-tree databases to investigate the intergenerational transmission of human longevity^22–25^. We concluded that longevity, as a heritable trait, is primarily transmitted if persons belong to the oldest 10% survivors of their birth cohort and if at least 30% of their ancestors also belonged to the oldest 10% survivors^22,23^. Subsequently, we developed the Longevity Relatives Count (LRC) score as an instrument to quantify the number of long-lived family members and observed that the survival advantage of study participants increased with each additional long-lived family member, indicating additive effects^22^. As such, the LRC score is an indicator of increased survival and longevity, and can therefore be used to enlarge the survival contrast in epidemiological data, thereby leading to more powerful genetic longevity studies. If the LRC score also represents healthspan as a quantitative trait (additive effects), this instrument can potentially be used in (genetic) studies to elucidate multi-morbidity limiting mechanisms.

We identified two issues that have not yet thoroughly been investigated: (1) whether from mid-life onward, health, medication use, disease incidence as well as the development of multi-morbidity are delayed over time, and (2) whether an increasing number of long-lived ancestors, as measured with the LRC score, represents not only lifespan as a quantitative trait but also healthspan. To address these issues, longitudinal life course and health data should ideally be investigated, preferably in large numbers of individuals. In addition, multiple generational family-tree information is required to investigate how the number of ancestral long-lived relatives relates to morbidity. Therefore we investigate chronic disease incidence and multimorbidity in long-lived families using up to 25 years of follow-up data. We further study whether an increasing number of long-lived ancestors, as measured by the LRC score, associates with a decreased incidence of chronic diseases. In addition, we investigate whether the families with the most extreme LRC scores have a healthy metabolomic profile in mid-life, representing overall health to complement the information on morbidity.

We use the data available in the Leiden Longevity Study (LLS, Netherlands) and the Swedish register data available in the Scanian Economic-Demographic Database (SEDD, Sweden). The LLS, initiated in 2002, was based on the inclusion of nonagenarian siblings. Also the middle aged children (called index persons (IPs) in the current study) and their partners, as adult environment-matched controls, were included. The SEDD contains the entire population of 5 parishes and a town in Scania (southern Sweden), and as such does not contain any initial inclusion criteria. For the current study we identified in both datasets combined 2,143 three-generational families (F1-F3) containing IPs (F3) and their family members, comprising 17,539 persons in total. First, we examine whether LLS IPs and their partners differ in terms of disease and medication prevalence at the moment of study inclusion (2002-2004). Second, we investigate differences in disease incidence towards multimorbidity. Third, we study whether an increasing number of long-lived ancestors is associated with a decreased disease incidence in IPs using the Longevity Relatives Count (LRC) score^23^ in both LLS and SEDD datasets. Finally, we compare mid-life health of LLS IPs with the highest LRC scores and their partners, using a previously developed metabolomics based score predicting mortality^26^.

## Results

### Study populations

LLS IPs and their partners, serving as environment-matched controls, were included between 2002 and 2006 at the average age of 59 years. The study inclusion was based on nonagenarian siblings in the F2 generation. Hence, IPs (F3) were included if they had at least one long-lived F2 parent and F2 aunt or uncle (females ≥ 91 years and males ≥ 89 years). From inclusion onward, the IPs and their partners were followed over time, with a maximum mortality follow-up of 19 years (2002-2021) and maximum morbidity follow-up of 16 years (2002-2018). In 2021, 227 (14%) IPs and 113 (15%) partners were deceased and 1409 (84%) IPs and 619 (83%) partners were still alive. In 2018, 671 (40%) IPs and 324 (43%) partners had a disease diagnosis whereas 535 (32%) IPs and 206 (28%) partners did not have a disease diagnosis (Figure 1A and Table 1A).

**Figure 1:**
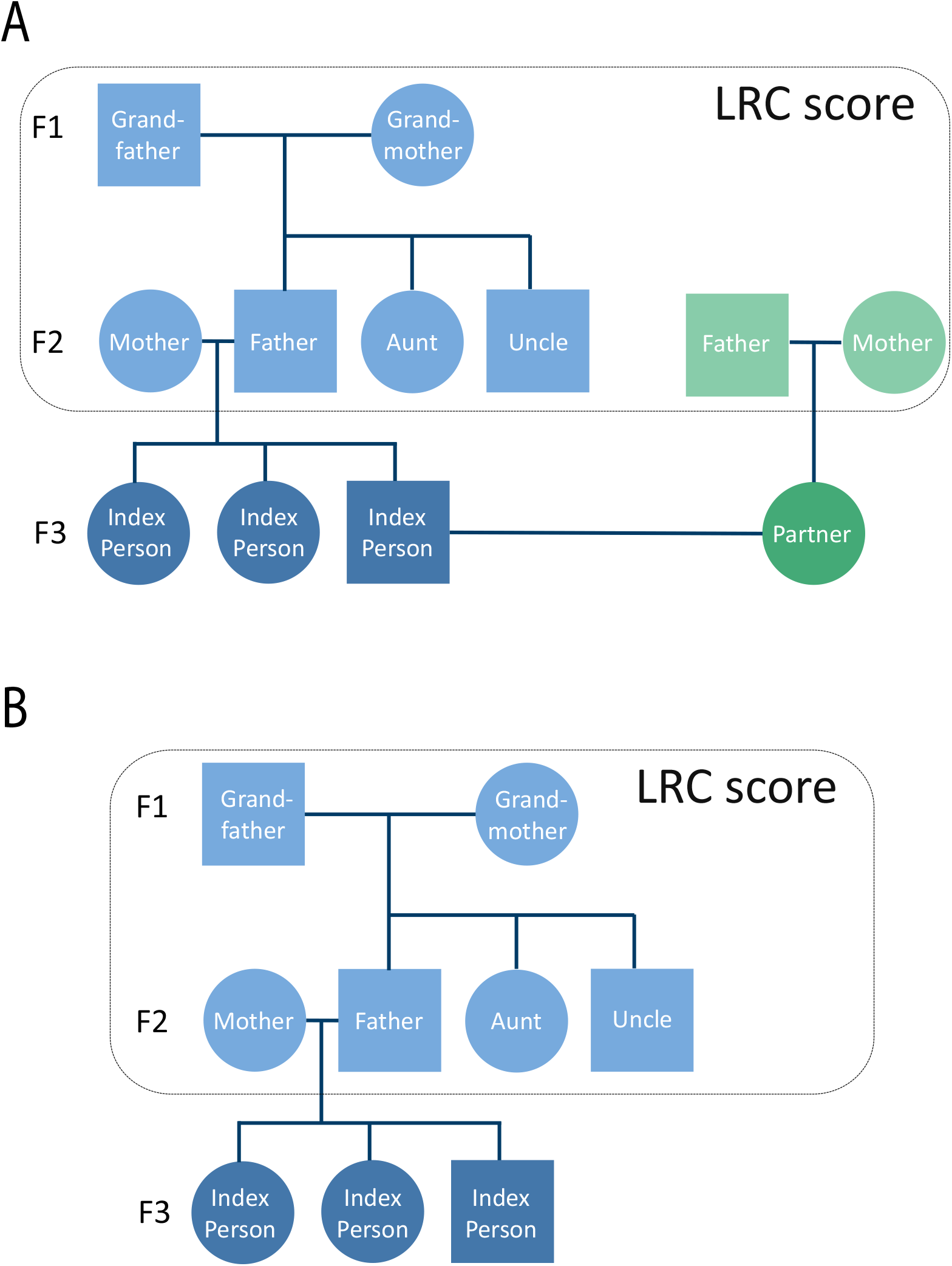
Conceptual pedigree of a 3 filial (F) generation LLS family. This figure corresponds to Table 1 and represents a hypothetical family from the LLS covering 3 filial (F) generations. Circles represent women, Squares represent men. Dark blue: Index persons (IPs, F3), dark green: partners of IPs (F3), light blue: fathers and mothers of IPs (F2), aunts and uncles of IPs (F2), grandmothers and grandfathers of IPs (f1), light green: fathers and mothers of IPs (F2). The dark blue and green colors represent the IPs and their partners who are investigated in this study. The light blue and green colors represent the ancestors of the IPs and partners and were used in this study to calculate the Longevity Relatives Count (LRC) score.

**Table 1:** Basic characteristics of LLS Index Persons, partners, and ancestral groups.

|  | <u>Analyses groups</u> |  | <u>Ancestral family groups</u> |  |  |  |
| --- | --- | --- | --- | --- | --- | --- |
|  | IPs F3 | Partners of IPs F3 | Parents of IPs F2 | Aunts/uncles of IPs F2 | Grandparents of IPs F1 | Parents of partners F2 |
| **<u>Panel A: LLS</u>** |  |  |  |  |  |  |
| Number, N | 1674 | 745 | 1295 | 2370 | 760 | 1237 |
| Female, N (%) | 905 (54) | 429 (58) | 646 (50) | 1169 (49) | 380 (5) | 620 (5) |
| Range birth cohorts, years | 1923-1971 | 1924-1974 | 1882-1928 | 1875-1951 | 1850-1894 | 1864-1947 |
| Alive, N (%) | 1409 (84) | 619 (83) | 116 (9) | 368 (16) | 0 (0) | 254 (21) |
| Deceased, N (%) | 227 (14) | 113 (15) | 1179 (91) | 2001 (84) | 760 (100) | 1011 (82) |
| Missing age, N (%) | 38 (2) | 13 (2) | 94 (7) | 34 (1) | 0 (0) | 35 (3) |
| Mean ad or al, years (SD) | 74.87 (6.67) | 74.21 (7.22) | 87.09 (14.13) | 71.93 (28.03) | 77.06 (14.15) | 76.68 (13.86) |
| Disease free, N (%) | 535 (32) | 206 (28) | - | - | - | - |
| Diseased, N (%) | 671 (40) | 324 (43) | - | - | - | - |
| Missing disease, N (%) | 468 (28) | 215 (29) | - | - | - | - |
| Mean number of diseases, N, (SD) | 0.82 (0.92) | 0.94 (0.98) | - | - | - | - |
| **<u>Panel B: SEDD</u>** |  |  |  |  |  |  |
| Number, N | 2497 | - | 2969 | 5830 | 3028 | - |
| Female, N (%) | 1252 (50.1) | - | 1480 (49.8) | 2866 (49.2) | 1532 (50.6) | - |
| Range birth cohorts, years | 1930-1950 | - | 1853-1914 | 1860-1950 | 1833-1918 | - |
| Alive, N (%) | 1803 (72.2) | - | 7 (0.2) | 855 (14.7) | 131 (4.3) | - |
| Deceased, N (%) | 694 (27.8) | - | 2962 (99.8) | 4975 (85.3) | 2897 (95.7) | - |
| Missing age, N (%) | - | - | - | - | - | - |
| Mean ad or al, years (SD) | 74.6 (6.7) | - | 76.0 (17.9) | 57.4 (32.2) | 64.0 (19.6) | - |
| Disease free, N (%) | 1307 | - | - | - | - | - |
| Diseased, N (%) | 1190 | - | - | - | - | - |
| Missing disease, N (%) | - | - | - | - | - | - |
| Mean number of diseases, N, (SD) | - | - | - | - | - | - |

SEDD IPs were followed over time from 1990, at an average age of 52 years, with a maximum mortality and morbidity follow-up of 25 years (1990-2015). In 2015, 694 (28%) IPs were deceased whereas 1,803 (72%) were still alive. Moreover, 1,190 (48%) IPs had a disease diagnosis whereas 1,307 (52%) IPs did not have a disease diagnosis (Figure 1B and Table 1B). From here we will refer to disease diagnoses as diseases, disease prevalence in cross-sectional analyses, and disease incidence in longitudinal analyses.

### Disease prevalence at LLS study inclusion

To investigate whether LLS IPs and their partners differ in terms of disease and medication prevalence at the moment of study inclusion we conducted mixed-model logistic regression analysis. Figure 2A shows a 13% lower odds for age-related disease prevalence in IPs (OR=0.87 (95% CI=0.64−1.17)) compared to their partners. We further observed that IPs had a 20% lower odds for metabolic diseases (OR=0.80 (95% CI=0.58−1.15)) and 16% higher odds for malignant diseases (OR=1.16 (95% CI=0.64−2.11) compared to their partners, albeit not statistically significant.

**Figure 2:**
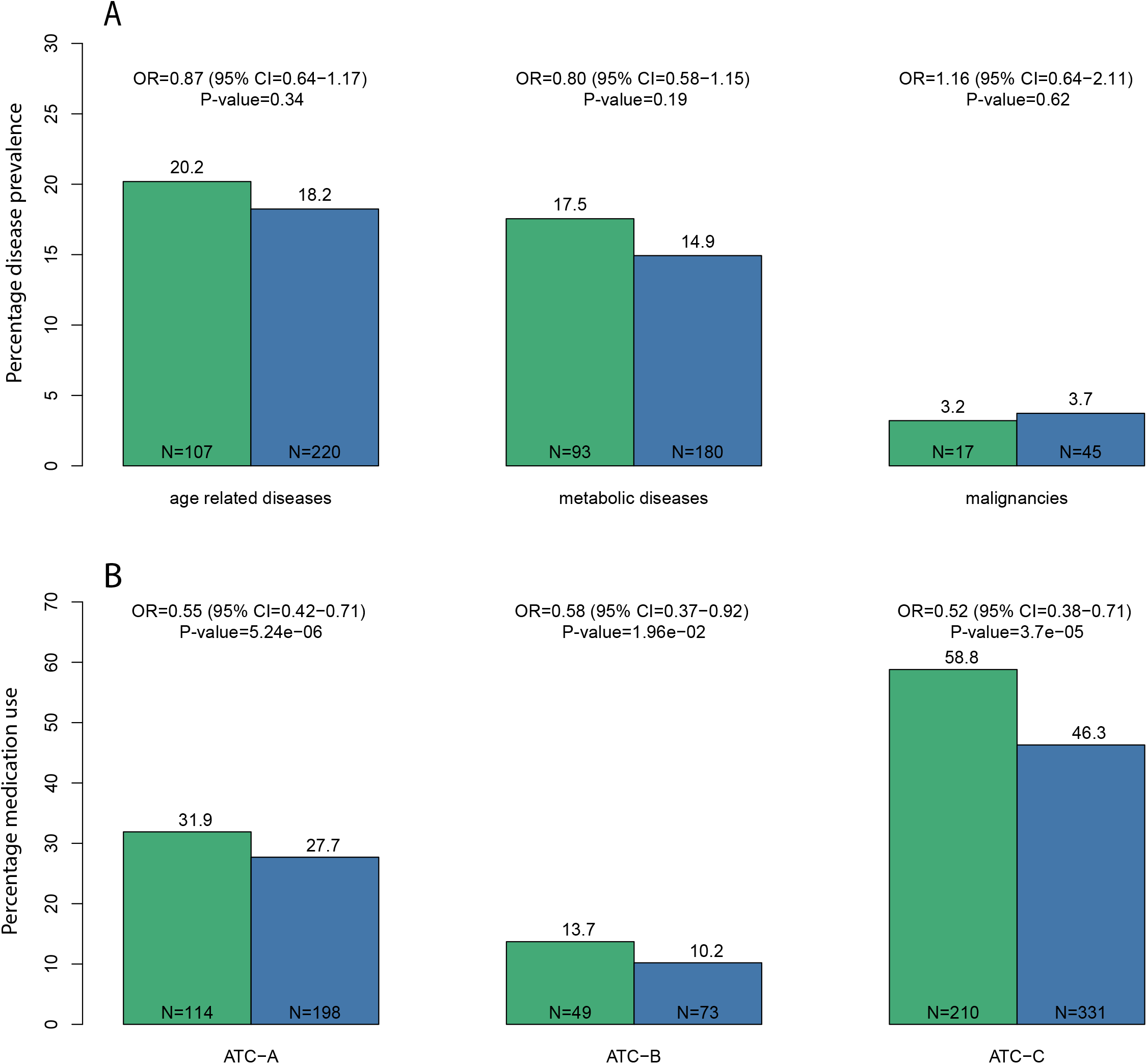
Disease prevalence and medication use in the LLS. This figure depicts the odds ratio’s (ORs) for disease prevalence (panel A) and medication use (Panel B). Blue bars represent LLS IPs and green bars represent their partners, similar to the colors used in Figure 1. The y-axis of panel A represent the percentage of LLS IPs and partners who had an age-related, metabolic, or malignant disease (x-axis). The y-axis of panel B represent the percentage of LLS IPs and partners who used ATC-A (alimentary tract and metabolism), ATC-B (blood and blood forming organs), or ATC-C (cardiovascular system) type medications (x-axis). CI is the abbreviation for confidence interval and N represents the numbers of the LLS IPs and partners in the specific disease groups. All estimates are adjusted for age at inclusion and sex.

### LLS IPs have a lower risk of using medication early in the study

Data on medication use was collected in the LLS between 2006 and 2008 (Supplementary Figure 1) and was available for 1254 LLS IPs (75%) and 588 partners (79%). We focused on ATC A-C type medication because they match the disease groups we investigate (see methods). To study whether LLS IPs had a lower medication use compared to their partners, we fitted mixed-model logistic regression analyses. Figure 2B shows that the odds of using ATC-A (alimentary tract and metabolism) type medication is 45% (OR=0.55 (95% CI=0.42−0.71)) lower for the offspring than for their partners. Similarly, the odds of using ATC-B (blood and blood forming organs) and ATC-C (cardiovascular system) type medication is 42% (OR=0.58 (95% CI=0.37−0.92)) and 48% (OR=0.52 (95% CI=0.38−0.71)) lower for the IPs. Our analyses thus indicate that, early on in the study, the IPs already had a significantly lower intake of metabolic and cardiovascular disease medication.

### LLS IPs have a delayed first disease onset during follow-up

To investigate whether and to what extent the onset of first disease was delayed for the LLS IPs compared to their partners during 16 years of follow-up, we excluded persons who had ≥1 disease at inclusion. We therefore include 917 LLS IPs of whom 39 (4.3%) were deceased at the end of disease follow-up (2018) and 395 partners of whom 17 (4.3%) were deceased. We fitted random effect (frailty) Cox regressions and observed a Hazard Ratio (HR) of 0.79 (95% CI=0.65-0.97) for the age-related disease incidence between LLS IPS and their partners. This HR indicates that the yearly risk of age-related disease was 21% lower for the LLS IPs as compared to their partners. The LLS IPs had a 29% (HR=0.71 (95% CI=0.55-0.90)) lower risk of metabolic diseases and a 5% (HR=0.95 (95% CI=(0.70-1.31)) lower risk of malignant diseases (Table 2A and Supplementary Table 1A). In addition, Supplementary Figure 2 shows that 50% of the LLS IPs had an age-related disease at the age of 68 years whereas this was the case at the earlier age of 65.8 years for their partners. 50% of the LLS IPs had a metabolic disease at an age of 74.8 years, while this was the case at 68.6 years for their partners, indicating a median delay of metabolic disease diagnosis for LLS IPs of 6.2 years.

**Table 2:** LLS disease incidence.

|  | N (prop) | Events (prop) | HR (95% CI) | P-Value |
| --- | --- | --- | --- | --- |
| A: Time from inclusion to first disease |  |  |  |  |
| Age related diseases |  |  |  |  |
| LLS IPs | 917 (0.70) | 362 (0.39) | 0.79 (0.65-0.97) | 2.32·10<sup>-2</sup> |
| Partners (ref) | 395 (0.30) | 171 (0.43) |  |  |
| Metabolic diseases |  |  |  |  |
| LLS IPs | 917 (0.70) | 261 (0.23) | 0.71 (0.55-0.90) | 5.18·10<sup>-3</sup> |
| Partners (ref) | 395 (0.30) | 135 (0.34) |  |  |
| Malignancies |  |  |  |  |
| LLS IPs | 917 (0.70) | 130 (0.14) | 0.95 (0.70-1.31) | 7.66·10<sup>-1</sup> |
| Partners (ref) | 395 (0.30) | 56 (0.14) |  |  |
| B: Time from inclusion to 2 diseases |  |  |  |  |
| Age related diseases |  |  |  |  |
| LLS IPs | 611 (0.70) | 73 (0.12) | 0.55 (0.36-0.85) | 6.83·10<sup>-3</sup> |
| Partners (ref) | 268 (0.30) | 47 (0.18) |  |  |
| Metabolic diseases |  |  |  |  |
| LLS IPs | 611 (0.70) | 45 (0.07) | 0.51 (0.26-0.97) | 3.96·10<sup>-2</sup> |
| Partners (ref) | 268 (0.30) | 29 (0.11) |  |  |
| Malignancies |  |  |  |  |
| LLS IPs | 611 (0.70) | 7 (0.01) | 1.39 (0.29-6.70) | 6.82·10<sup>-1</sup> |
| Partners (ref) | 268 (0.30) | 2 (0.01) |  |  |
| C: Time from first disease to second disease |  |  |  |  |
| Age related → age related diseases |  |  |  |  |
| LLS IPs | 500 (0.68) | 79 (0.16) | 0.46 (0.26-0.83) | 9.82·10<sup>-3</sup> |
| Partners (ref) | 237 (0.32) | 55 (0.23) |  |  |
| Age related → Metabolic diseases |  |  |  |  |
| LLS IPs | 500 (0.68) | 62 (0.12) | 0.33 (0.14-0.81) | 1.46·10<sup>-2</sup> |
| Partners (ref) | 237 (0.32) | 44 (0.19) |  |  |
| Age related → Malignant diseases |  |  |  |  |
| LLS IPs | 500 (0.68) | 17 (0.03) | 0.58 (0.27-1.25) | 1.66·10<sup>-1</sup> |
| Partners (ref) | 237 (0.32) | 11 (0.05) |  |  |

### LLS IPs have a delayed onset of multimorbidity during follow-up

To study whether the delayed onset of first disease for LLS IPs extended to developing more than one disease (multimorbidity) during the 16 years of follow-up, we investigated the difference in time from inclusion to having two diseases within the same category (2 age-related, metabolic, or malignant diseases; Table 2B and Supplementary Table 1B). We observed that the yearly risk to develop 2 age-related diseases was 45% (HR=0.55 (95% CI=0.36-0.85) lower for the LLS IPs than for their partners, maximizing to a 49% (HR=0.51 (95% CI=0.26-0.97) difference for metabolic diseases. However, the yearly risk to develop 2 malignant diseases (HR=1.39 (95% CI=0.29-6.70)) did not significantly differ between LLS IPs and their partners. Supplementary Figure 3 shows the survival curves corresponding to Table 2B.

To study whether LLS IPs, who already had a disease, had a lower risk of getting a second disease, we investigated whether the time between first and specific second disease was longer for the LLS IPs than for their partners. Table 2C and Supplementary Table 2 show that the yearly risk to develop an age-related or a metabolic disease after already being diagnosed with an age-related disease, was 54% (HR=0.46 (95% CI=0.26-0.83), and 66% (HR=0.33 (95% CI=0.14-0.81)) lower for the IPs, respectively. For a malignant disease following the initial diagnosis of an age-related disease, no significant difference between LLS IPs and their partners was observed (HR=0.58 (95% CI=0.27-1.25)). Supplementary Figure 4 shows the survival curves corresponding to Table 2C. Moreover, sensitivity analyses showed that the HRs representing time from first to second disease were not affected by the group (metabolic or malignant) of first disease.

### Increasing numbers of long-lived ancestors indicate a later disease onset in LLS and SEDD

In our previous work we developed the Longevity Relatives Count (LRC) score to quantify a person’s number of long-lived ancestors^22,23^. The LRC score can be interpreted as a weighted proportion (ranging between 0 and 1)^23^. For example, an LRC score of 0.5 for an IP indicates 50% long-lived ancestors weighted by the genetic distance between IPs (and partners in LLS) and their ancestors. Here we investigate whether healthspan in LLS and SEDD is associated with the number of long-lived ancestors by testing whether an increasing LRC score of IPs is associated with a delay in disease onset and lower medication use in a longitudinal study design of the two independent databases; LLS and SEDD.

We conducted our analyses using two approaches. In the first approach we used the LRC score to enlarge the contrasts between the LLS IPs and their partners by defining four mutually exclusive groups: LRC_g1: IPs with an LRC ≥0.60, LRC_g2: IPs with an LRC [≥0.1 & <0.60], LRC_g3: partners with an LRC >0, and LRC_g4: partners with an LRC =0. We subsequently compared the disease incidence and medication use of LRC_g1-3 with LRC_g4, using Cox-type random effect (frailty) and linear mixed model regression analysis respectively. In the second approach, we calculated the LRC score in the LLS IPs and partners combined, allowing a quantitative definition of the LRC-score instead of defining groups. Using the quantitatively defined LRC-score we investigate whether an increasing LRC score associates with a decreasing first disease incidence, using Cox-type random effect (frailty) regression analysis. Finally, we validate the results obtained in the LLS by replicating our analysis in Swedish register data (SEDD).

#### First approach

When comparing the LLS IPs with an LRC score ≥0.60 (LRC_g1) with the partners who had an LRC score of 0 (LRC_g4) we observed a HR of 0.56 (95% CI=0.34-0.92) and 0.69 (95% CI=0.31-1.53) for time to first age-related and malignant disease, respectively (Table 3). Table 3 further shows that the healthspan benefit of the LRC_g1 group was most striking for the incidence of first metabolic disease, for which the yearly risk was 53% lower (HR=0.47 (95% CI=0.25-0.87)). For comparison: HR’s in Table 2 (not applying LRC score) are 0.79, 0.95 and 0.71 for age-related, malignant and metabolic diseases respectively, providing a strong indication that increasing numbers of long-lived ancestors are associated to a later disease incidence. To illustrate this comparison, Figure 3 shows the survival curves for the LLS IPs and partners (Panel A corresponding to Table 2) and the LRC groups (Panel B corresponding to Table 3). The figure shows how the LRC score maximizes the contrast: 50% of the LRC_g4 persons developed a first metabolic disease at the age of 68 years, whereas 50% of the LRC_g1 persons developed a first metabolic disease at the age of 81 years. Hence, the LRC_g1 persons delayed the age of metabolic disease onset with a pronounced 13 years difference. The survival curves of the other disease categories are presented in Supplementary Figure 5. Further benefit for LRC_g1 over LRC_g4 concerns development of multimorbidity and medication use, since an HR of 0.14 (95% CI=0.03-0.70) was observed for the time to develop 2 age-related diseases and an OR of 0.26 (95% CI=0.12-0.57) for medication use.

**Table 3:**
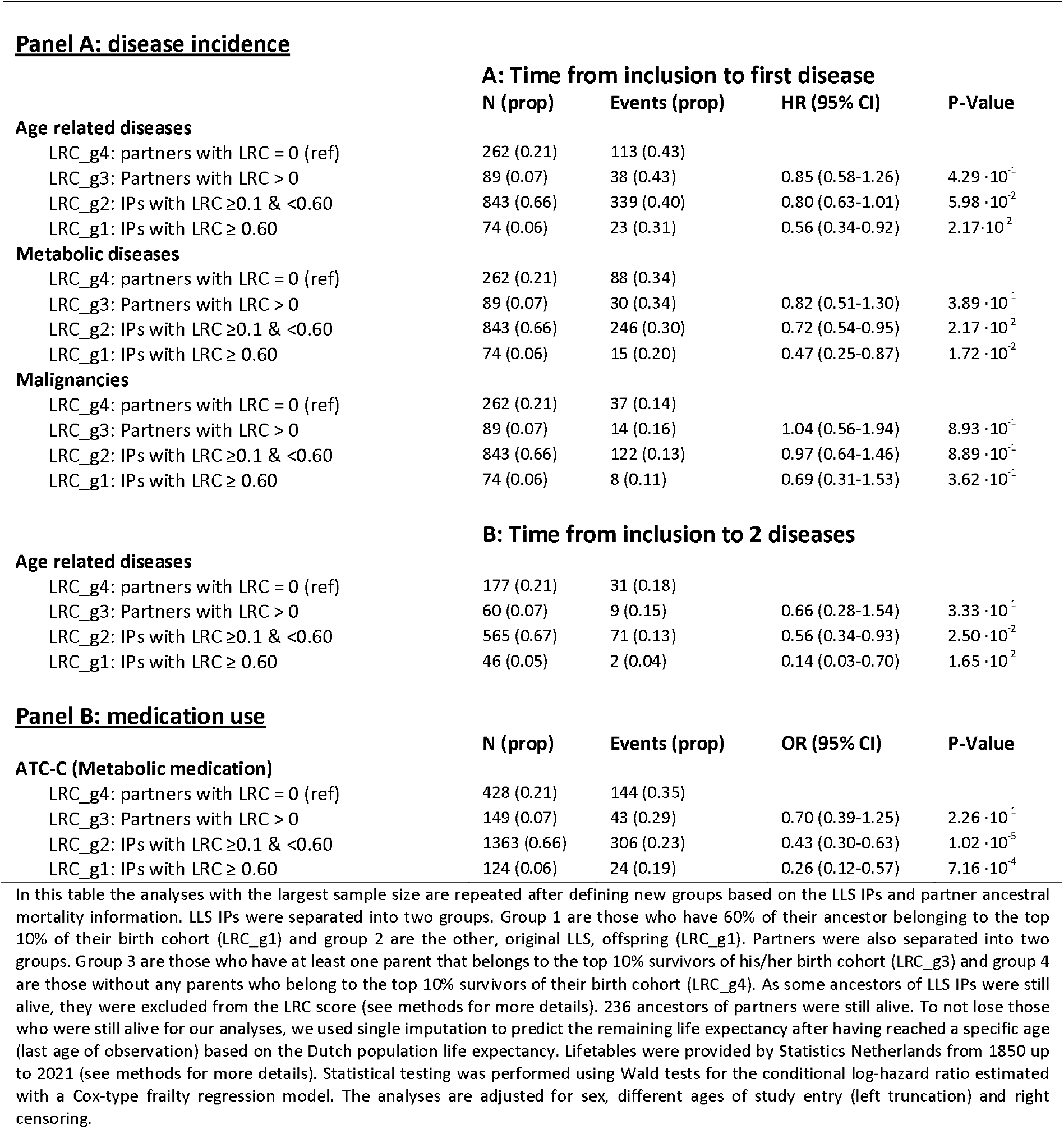
LLS disease incidence and medication use in LLS LRC groups.

**Panel A: disease incidence**
|  | **A: Time from inclusion to first disease** |  |  |  |
| --- | --- | --- | --- | --- |
|  | **N (prop)** | **Events (prop)** | **HR (95% CI)** | **P-Value** |
| **Age related diseases** |  |  |  |  |
| LRC_g4: partners with LRC = 0 (ref) | 262 (0.21) | 113 (0.43) |  |  |
| LRC_g3: Partners with LRC > 0 | 89 (0.07) | 38 (0.43) | 0.85 (0.58-1.26) | $4.29 \cdot 10^{-1}$ |
| LRC_g2: IPs with LRC $\geq 0.1$ & <0.60 | 843 (0.66) | 339 (0.40) | 0.80 (0.63-1.01) | $5.98 \cdot 10^{-2}$ |
| LRC_g1: IPs with LRC $\geq 0.60$ | 74 (0.06) | 23 (0.31) | 0.56 (0.34-0.92) | $2.17 \cdot 10^{-2}$ |
| **Metabolic diseases** |  |  |  |  |
| LRC_g4: partners with LRC = 0 (ref) | 262 (0.21) | 88 (0.34) |  |  |
| LRC_g3: Partners with LRC > 0 | 89 (0.07) | 30 (0.34) | 0.82 (0.51-1.30) | $3.89 \cdot 10^{-1}$ |
| LRC_g2: IPs with LRC $\geq 0.1$ & <0.60 | 843 (0.66) | 246 (0.30) | 0.72 (0.54-0.95) | $2.17 \cdot 10^{-2}$ |
| LRC_g1: IPs with LRC $\geq 0.60$ | 74 (0.06) | 15 (0.20) | 0.47 (0.25-0.87) | $1.72 \cdot 10^{-2}$ |
| **Malignancies** |  |  |  |  |
| LRC_g4: partners with LRC = 0 (ref) | 262 (0.21) | 37 (0.14) |  |  |
| LRC_g3: Partners with LRC > 0 | 89 (0.07) | 14 (0.16) | 1.04 (0.56-1.94) | $8.93 \cdot 10^{-1}$ |
| LRC_g2: IPs with LRC $\geq 0.1$ & <0.60 | 843 (0.66) | 122 (0.13) | 0.97 (0.64-1.46) | $8.89 \cdot 10^{-1}$ |
| LRC_g1: IPs with LRC $\geq 0.60$ | 74 (0.06) | 8 (0.11) | 0.69 (0.31-1.53) | $3.62 \cdot 10^{-1}$ |
| **B: Time from inclusion to 2 diseases** |  |  |  |  |
| **Age related diseases** |  |  |  |  |
| LRC_g4: partners with LRC = 0 (ref) | 177 (0.21) | 31 (0.18) |  |  |
| LRC_g3: Partners with LRC > 0 | 60 (0.07) | 9 (0.15) | 0.66 (0.28-1.54) | $3.33 \cdot 10^{-1}$ |
| LRC_g2: IPs with LRC $\geq 0.1$ & <0.60 | 565 (0.67) | 71 (0.13) | 0.56 (0.34-0.93) | $2.50 \cdot 10^{-2}$ |
| LRC_g1: IPs with LRC $\geq 0.60$ | 46 (0.05) | 2 (0.04) | 0.14 (0.03-0.70) | $1.65 \cdot 10^{-2}$ |

**Panel B: medication use**
|  | **N (prop)** | **Events (prop)** | **OR (95% CI)** | **P-Value** |
| --- | --- | --- | --- | --- |
| **ATC-C (Metabolic medication)** |  |  |  |  |
| LRC_g4: partners with LRC = 0 (ref) | 428 (0.21) | 144 (0.35) |  |  |
| LRC_g3: Partners with LRC > 0 | 149 (0.07) | 43 (0.29) | 0.70 (0.39-1.25) | $2.26 \cdot 10^{-1}$ |
| LRC_g2: IPs with LRC $\geq 0.1$ & <0.60 | 1363 (0.66) | 306 (0.23) | 0.43 (0.30-0.63) | $1.02 \cdot 10^{-5}$ |
| LRC_g1: IPs with LRC $\geq 0.60$ | 124 (0.06) | 24 (0.19) | 0.26 (0.12-0.57) | $7.16 \cdot 10^{-4}$ |

In this table the analyses with the largest sample size are repeated after defining new groups based on the LLS IPs and partner ancestral mortality information. LLS IPs were separated into two groups. Group 1 are those who have 60% of their ancestor belonging to the top 10% of their birth cohort (LRC\_g1) and group 2 are the other, original LLS, offspring (LRC\_g1). Partners were also separated into two groups. Group 3 are those who have at least one parent that belongs to the top 10% survivors of his/her birth cohort (LRC\_g3) and group 4 are those without any parents who belong to the top 10% survivors of their birth cohort (LRC\_g4). As some ancestors of LLS IPs were still alive, they were excluded from the LRC score (see methods for more details). 236 ancestors of partners were still alive. To not lose those who were still alive for our analyses, we used single imputation to predict the remaining life expectancy after having reached a specific age (last age of observation) based on the Dutch population life expectancy. Lifetables were provided by Statistics Netherlands from 1850 up to 2021 (see methods for more details). Statistical testing was performed using Wald tests for the conditional log-hazard ratio estimated with a Cox-type frailty regression model. The analyses are adjusted for sex, different ages of study entry (left truncation) and right censoring.

**Figure 3:**
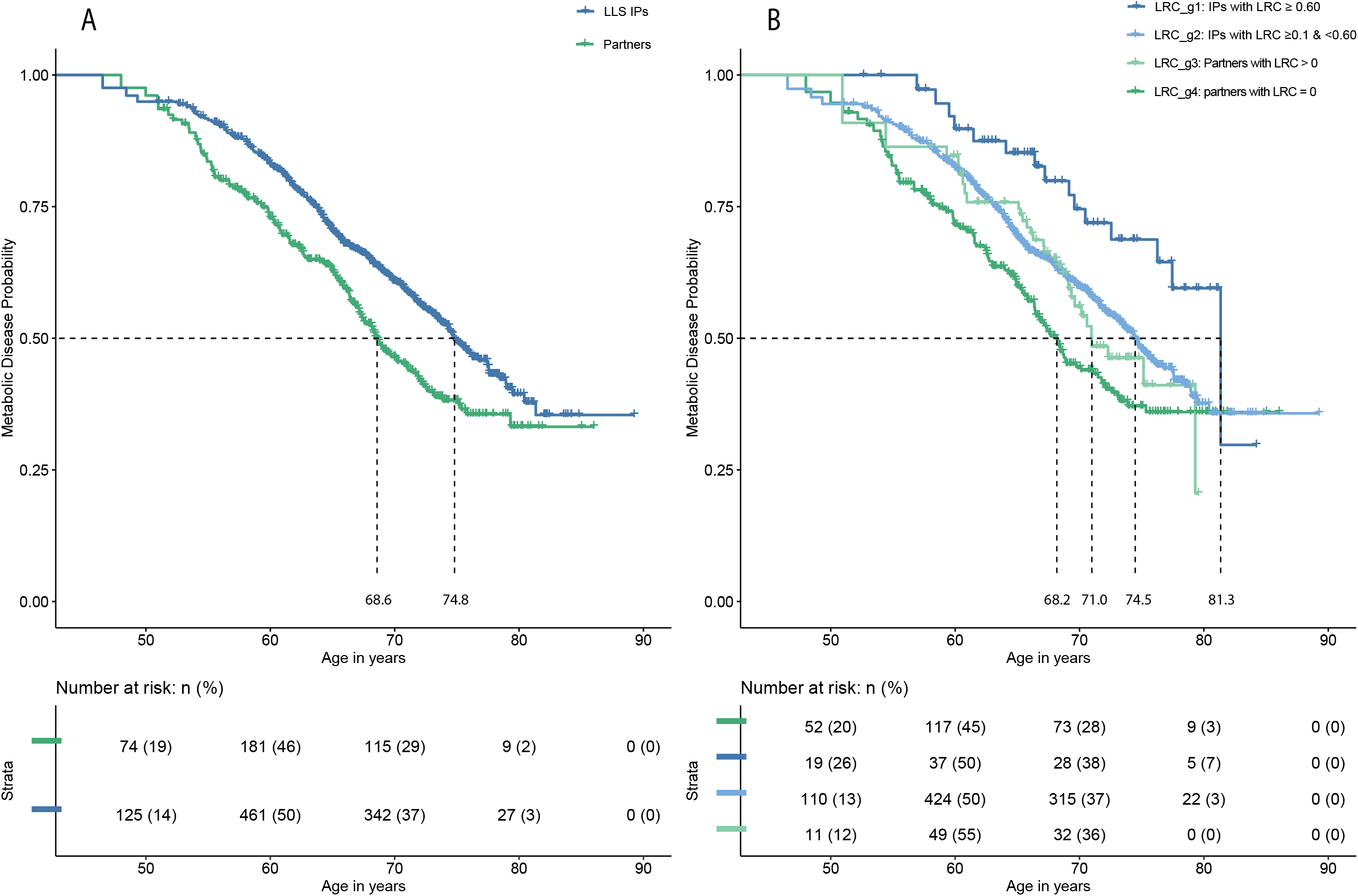
LLS metabolic disease incidence with and without LRC-defined groups. This figure depicts survival curves reflecting metabolic disease incidence within the Leiden Longevity Study (LLS). The x-axis show age in years and the y-axis show metabolic disease incidence. Dotted lines represent the age at which 50% of the members of a specific group had their first metabolic disease. Panel A depicts two groups; the blue line represents LLS Index Persons (IPs) and the green line represents the partners. The mean difference between the lines represents the Hazard Ratio (HR) shown in Table 2. Panel B depicts four groups; LRC_g1: IPs with an LRC ≥0.60 (dark blue), LRC_g2: IPs with an LRC [≥0.1 & <0.60] (light blue), LRC_g3: partners with an LRC >0 (light green), and LRC_g4: partners with an LRC=0 (dark green). The mean difference between the LRC_g1-3 and LRC_g4 line represents the HR shown in Table 3. Vertical lines within the colored lines represent right censoring events. The bottom column of panel A and B shows how many persons were still at risk of having a metabolic disease at different ages. Survival curves are adjusted for left truncation and right censoring.

#### Second approach

We calculated the LRC score for the LLS IPs and their partners combined to avoid any grouping. Table 4A shows that with every 0.1 (10%) increase in LRC score, LLS F3 participants had a 5% (HR=0.95 (95% CI=0.91-0.99)) lower yearly risk to develop a first age-related disease. To illustrate the magnitude, this effect increases to 50% when all ancestors were long-lived (LRC score = 1). We further observed a 7% (HR=0.93 (95% CI=0.88-0.98)) lower yearly first metabolic and 3% (HR=0.97 (0.91-1.04)) lower malignant disease risk, though the latter effect was not statistically significant.

**Table 4:** Quantitative LRC analyses of time to first disease in LLS en SEDD.

|  | N | Events (prop) | HR (95% CI) | P-Value |
| --- | --- | --- | --- | --- |
| **A: LLS data** |  |  |  |  |
| Age related diseases | 1312 | 533 (0.41) | 0.95 (0.91-0.99) | $3.06 \cdot 10^{-2}$ |
| Metabolic diseases | 1312 | 396 (0.30) | 0.93 (0.88-0.98) | $1.37 \cdot 10^{-2}$ |
| Malignancies | 1312 | 186 (0.14) | 0.97 (0.91-1.04) | $4.98 \cdot 10^{-1}$ |
| **B: SEDD data** |  |  |  |  |
| Age related diseases | 2497 | 1,190 (0.48) | 0.94 (0.89-0.98) | $1.22 \cdot 10^{-3}$ |
| Metabolic diseases | 2497 | 706 (0.28) | 0.91 (0.87-0.96) | $4.84 \cdot 10^{-3}$ |
| Malignancies | 2497 | 671 (0.27) | 0.95 (0.90-0.99) | $4.43 \cdot 10^{-2}$ |
| Death | 2497 | 694 (0.28) | 0.92 (0.87-0.97) | $1.72 \cdot 10^{-3}$ |

Table shows the time from inclusion to first disease. N is the group size. Events are the events of the specific diseases, for example age related diseases, and prop. indicates the proportion from the size of a specific disease group. for example, 533 (41%) out of the 1312 persons had an age-related disease. sd indicates the standard deviation. HR is the abbreviation for Hazard Ratio. Statistical testing was performed using Wald tests for the conditional log-hazard ratio estimated with a Cox-type frailty regression model. The analyses are adjusted for sex, different ages of study entry (left truncation) and right censoring. Only persons without any disease at inclusion are studied.

We validate the results obtained in the LLS by replicating our analysis in Swedish register data (SEDD). Table 4B shows that with every 10% increase in LRC score, the SEDD IPs have a 6% (HR=0.94 (95% CI=0.89-0.98)) lower yearly risk to develop a first age-related disease, 9% (HR=0.91 (0.87-0.96)) lower risk for metabolic and 5% (HR=0.95 (0.90-0.99)) for malignant disease. Moreover, the yearly risk of dying decreases 8% (HR=0.92 (0.87-0.97)) with every 10% increase in LRC score.

### LLS IPs with an LRC score ≥0.60 already had a healthy metabolomic profile at inclusion

Our results point strongly towards protection from metabolic diseases for persons with an increasing number of long-lived ancestors as established with a high LRC score. We therefore investigated whether those with a high LRC score at the moment of inclusion in the LLS, indeed have a healthy circulating metabolomic profile that marks protection from disease at midlife. To estimate health in a quantitative parameter, we use a recently developed NMR-metabolomics based predictor of 5-10 years all-cause mortality across all ages from midlife onwards (from here MetaboHealth score)^26^. Hence, we explored whether the MetaboHealth score associates with differences between LRC groups as defined in the first approach of the analysis above (LRC_g1-3 compared to LRC_g4) and conducted a mixed model linear regression analysis.

We observed that the IPs with an LRC score ≥0.60 (LRC_g1 IPs) had a 0.098 (95%CI: [-0.184] – [-0.012]) lower MetaboHealth score than the partners who had an LRC score of 0 (LRC_g4 IPs; Figure 4 and Supplementary Table 3). The LRC_g2 and LRC_g3 IPs had a 0.032 (95%CI: [-0.077 - [0.012]) and a 0.016 (95%CI: [-0.091] - [0.058]) lower score than the LRC_g4 IPs respectively. Though the effects are relatively small (Figure 4), Indeed we observed that the LLS IPs with ≥60% long-lived ancestors who show delayed onset of disease, also have a healthier circulating metabolic profile in mid-life than the partners with an LRC score of 0.

**Figure 4:**
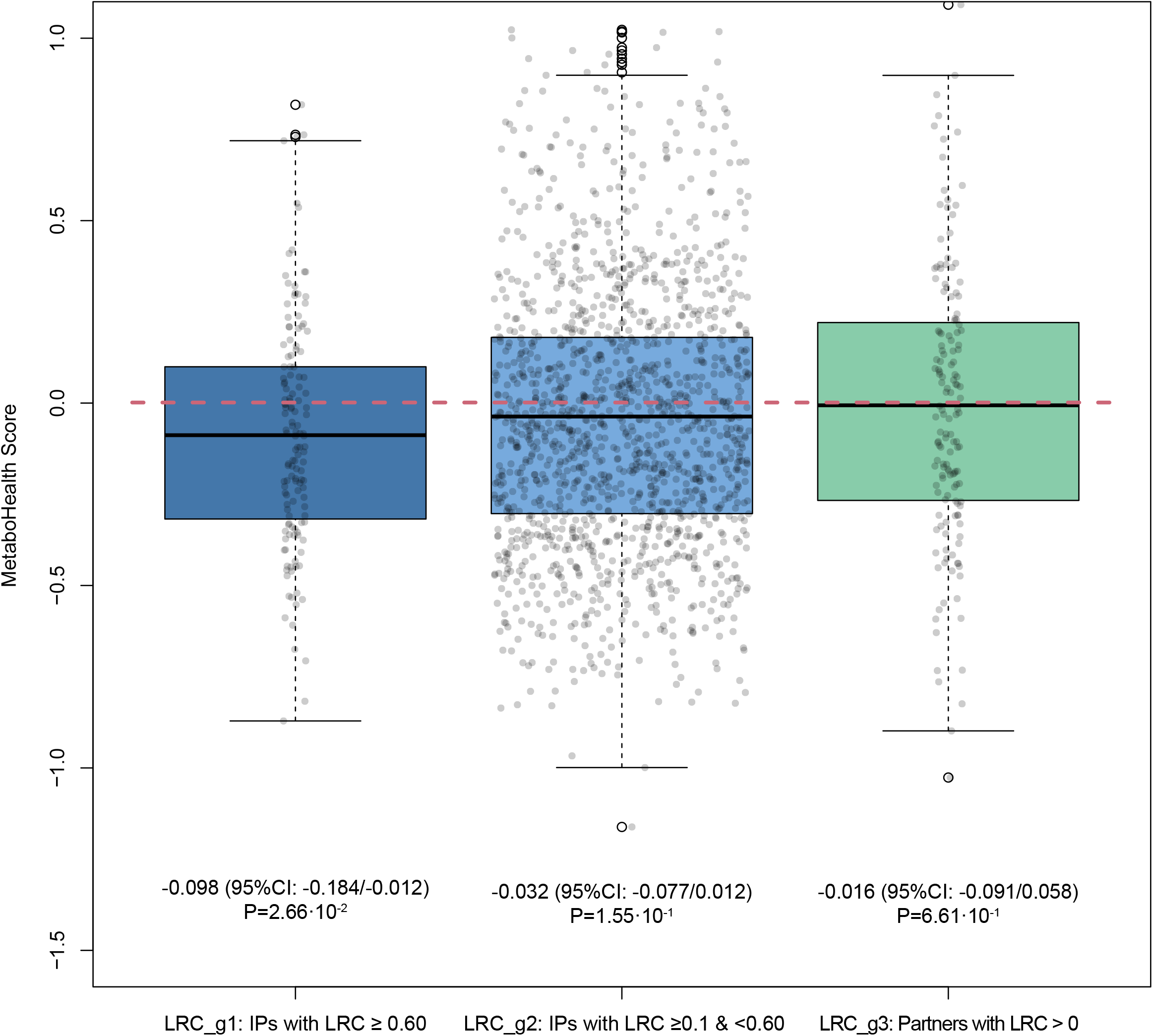
MetaboHealth score differences for LRC groups at LLS study inclusion. The x-axis depict three groups; LRC_g1: IPs with an LRC ≥0.60 (dark blue), LRC_g2: IPs with an LRC [≥0.1 & <0.60] (light blue), and LRC_g3: partners with an LRC >0 (light green). The dotted red line depicts the LRC_g4 group: partners with an LRC =0 (dark green). The y-axis depict the MetaboHealth score. Higher MetaboHealth score values represent a less healthy metabolomic profile as measured by the MetaboHealth score which represents 5/10 year mortality risk (see methods for more details). Error bars represent confidence intervals. CI is the abbreviation for confidence interval.

## Discussion

Human longevity is heritable and clusters in specific families. Members of these families live longer and seem to age healthier than the general population. Studying these long-lived families is important to improve our understanding of the molecular and environmental mechanisms that protect from (multi)morbidity and promote a healthy survival up to high ages. In this study we investigated the development of diseases from mid-life onwards in big multigenerational and prospective data, covering up to 25 years of follow-up, in family based (LLS, Netherlands) and population based (SEDD, Sweden) data. We showed that members of long-lived families have a delayed onset of disease, multimorbidity and medication use as compared to their partners, thereby extending their healthspan with up to a decade. These members also postponed multimorbidity since those who were already diagnosed with an age-related disease had a 54% lower risk of having a second age-related disease compared to their partners. When defining familial longevity quantitatively using the LRC score, we demonstrated that an increasing number of long-lived ancestors associates with an increasing delay in disease onset and lower medication use. Finally we demonstrated that at the moment of LLS study inclusion, those with an LRC score ≥0.60 (LRC_g1)) had a better MetaboHealth score than their partners with an LRC score of 0 (LRC_g4), indicating better immune and metabolic health, and lower 5-years mortality risk. We conclude that an increasing number of long-lived ancestors, as measured with the LRC score, is a quantitative indicator of familial longevity, capturing delayed mortality, protection against multimorbidity, and improved health in selected families as well as the population at large. The LRC score can thus potentially be used in genetic studies to elucidate multi-morbidity limiting mechanisms that promote healthspan already in mid-life.

Our analyses, using ancestral mortality data, in the selected Dutch longevity families and the Swedish register data led to remarkably similar conclusions. An increasing number of long-lived ancestors, as measured with the LRC score, not only associates with a lower mortality at any moment in life^22,23^, it also associates, in a similar way, with a lower disease incidence during mid and later life (60 to 75 years): With every 10% increase in LRC score the yearly risk to develop an age-related disease decreased with 39% in the LLS, maximizing to 46% in the SEDD.

We did observe stronger effects in the SEDD data than in the LLS data, with consistently lower hazard ratio’s (HRs) for age-related and metabolic disease incidence. This may be explained firstly because LLS IPs are compared with their partners as controls, either as separate or combined groups in the LRC analyses. IPs and partners share the same adult household and thus, the LLS design controls for shared resources and behavior (such as socio-economic status, social network, and lifestyle). In the SEDD data we did not compare IPs with their partners. The effect size difference between LLS and SEDD may therefore represent the influence of shared resources and behavior. Secondly, in the LLS disease diagnoses were obtained from the general practitioners (GPs) whereas in the SEDD, disease diagnoses were obtained from hospital records available in the Swedish national register data. It may be that stronger effects were observed in the SEDD because hospitalization is on average an indication of more extreme health events than receiving a GP diagnosis. Nevertheless, for many of the GP diagnosed diseases, such as a myocardial infarction, hospitalization is also required.

We did not observe statistically significant results for malignancies in the LLS data whereas we did observe significant effects in the SEDD data. However, within the SEDD data the effects for malignant disease incidence were considerably smaller (HR closer to reference group) than for metabolic and age-related diseases. A first explanation for this observation relates to differences in study population and follow-up time. LLS IPs and partners were followed-up for a maximum of 16 years from the average age of 59 years whereas for the SEDD IPs this was a maximum of 25 years from the average age of 52 years. As a result, we may have missed early onset (around 50 years) malignancies in the LLS whereas those are included in the SEDD. This may also explain why a considerably lower proportion of malignant diseases was available in LLS than in the SEDD whereas this was not the case for metabolic diseases. Secondly, inherited genetic factors have a limited effect on many types of malignancies, with heritability estimates ranging between 20% and 30%^27^. However, the chronic diseases, as measured in our study, are much more heritable^28–37^, with over 70% heritability for type 2 diabetes^38,39^. As the LRC score captures additive genetic effects^22,23^, the lower heritability of malignancies could explain why the small number of malignant disease observations in the LLS did not provide enough power to detect effects and why the effect sizes are lower in the SEDD compared to metabolic diseases. Previous research focusing on malignant diseases in long-lived individuals and their offspring obtained very heterogenous results^4,5,7,8,10,11,13–15,17,40–44^ which may also be due to study population and selection differences.

Past research primarily focused on studying disease prevalence of long-lived individuals, such as centenarians, and their children^4,10–14,17–20^ in cross sectional designs. Our data covers up to 25 years of follow-up and provides a unique combination between ancestral mortality information and deep phenotyping of chronic age-related diseases, medication use, as well as metabolomics. This allowed us to closely link familial longevity to medication use and the incidence of multiple diseases. Detailed information about disease incidence was provided by the treating physicians (General Practitioners, GPs) of the LLS IPs and their partners. In the SEDD, we used hospitalization records from the Swedish national registers (see methods for more details). The combination between these two types of data ensured robustness against healthy participant bias. Apart from this, the LLS was initiated with the inclusion of (1) LLS IPs who had at least one long-lived parent and aunt or uncle and (2) the partners of the LLS IPs. LLS IPs are likely to share social (e.g. social network, socio-economic status) and behavioral (e.g. lifestyle, drinking, sporting) traits important for healthy aging and longevity, for example because they share the same household or due to assortative mating^45^. The LLS study design thus corrects for such similarities between LLS IPs and partners, potentially resulting in an underestimation of differences between LLS IPs and partners as compared to general population controls. However, replication in the SEDD, which does not contain any initial inclusion criteria, guarantees results which are not influenced by partner similarities.

Genetic longevity studies so far mainly focused on survival to exceptional ages. Using the LRC score, disease-free survival, possibly in combination with the MetaboHeath score, may be explored as a broader phenotype to increase the power of longevity genetic studies. In addition, the association between LRC score and delayed disease incidence may be explained by the presence of variants protecting from development of disease and/or the absence of disease loci in the long-lived families. Though, previous research showed no evidence that long-lived persons were characterized by the absence of disease loci^46^, GWAS studies identifying disease susceptibility variants for example, for hypertension^47^, Alzheimer’s disease^48^ has progressed significantly. Hence, it is interesting re-investigate if the absence of disease susceptibility loci associates to the LRC score. As mentioned in the previous paragraphs, the larger effect sizes in the SEDD likely illustrate the importance of shared resources and behavior in long-lived families. Further evidence for the clustering of socio-behavioral traits was provided in a recent study which showed that members of long-lived families were less frequently hospitalized with smoking-related cancers as a first disease^8^. As socio-behavioral traits are influenced by complex combinations between genes and environment, further investigation may aid genetic research while providing an interesting basis to investigate the social complexity underlying familial longevity.

Our results provide strong evidence that an increasing number of long-lived ancestors associates with up to a decade of healthspan extension and a healthy metabolomic profile in mid-life. Our results have two important implications. First, future genetic research aimed at identifying protective longevity mechanisms beneficially influencing the risk of multimorbidity could focus on a broader definition of longevity entailing survival to exceptional ages as well as disease free survival and possibly the MetaboHealth score metabolites. Second, our results highlight the importance of integrating multiple generations of ancestral mortality data to existing and novel studies. In the past it was difficult to obtain such ancestral information but currently it is much more feasible to do so, as population scale family tree data is becoming increasingly available^22,49,50^. Moreover, in an increasing number of countries ancestral data can be retrieved from the national statistics bureaus, such as Statistics Sweden or Statistics Netherlands. Finally, next to genetic drivers of longevity and disease incidence, it is important to investigate if and how potential socio-behavioral resources^51^, such as socio-economic status and stress, associate to both longevity and disease incidence. If these novel insights are consistently applied across studies, the comparative nature of longevity studies may improve and facilitate the discovery of novel genetic variants and mechanisms promoting healthy ageing.

## Methods

### Leiden Longevity Study

The Leiden Longevity Study (LLS) was initiated in 2002 to study the mechanisms that lead to exceptional survival. The LLS currently consist of 650 three-generational families, defined by siblings who have the same parents (Figure 1). Inclusion took place between 2002 and 2006 and initially started with the recruitment of living sibling pairs. Within a sibling pair, males were invited to participate if they were 89 years or older and females if they were 91 years or older. Inclusion was subsequently extended to the children of the sibling pairs and their partners. This study focuses on the children of the sibling pairs and their partners, referring to them as LLS IPs and partners. From their perspective, IPs were included if they had at least one long-lived parent and aunt or uncle (females ≥ 91 years and males ≥ 89 years). In total, 1,674 Index Persons (IPs, F3), 745 partners (F3), 1,295 parents (F2), 2,370 aunts and uncles (F2), 760 grandparents (F1), and 1,237 parents of the partners (F2) were included in this study.

Mortality information was verified by birth or marriage certificates and passports whenever possible. Additionally, verification took place via personal cards which were obtained from the Dutch Central Bureau of Genealogy. In January 2021 all mortality information was updated through the Personal Records Database (PRD) which is managed by Dutch governmental service for identity information. https://www.government.nl/topics/personal-data/personal-records-database-brp. The combination of officially documented information provides very reliable and complete ancestral as well as current mortality information.

Disease data has been retrieved from the General Practitioners (GPs) of the LLS IPs and partners and covers the period from birth until 2018. GPs extracted the presence of chronic age-related diseases as specified in Supplementary Table 4 and the year the disease occurred from their electronic health records. The GP records are kept up to date when a person switches from one GP practice to another. Diseases were clustered into 3 groups based on the International Statistical Classification of Diseases and Related Health Problems (ICD-10) codes, (1) metabolic diseases, (2) malignant diseases, and (3) age related diseases, which are the combination of metabolic and malignant diseases. Furthermore, cross-sectional information on medication use from pharmacies was obtained for the period 2006-2008, indicating whether a specific medicine was used or not. Medication was grouped according to the international Anatomical Therapeutic Chemical Classification System (ATC) standard. We focused on ATC-A (alimentary tract and metabolism), ATC-B (blood and blood forming organs), and ATC-C (cardiovascular system) type medications because they match the disease groups we investigate.

Ethylenediamine tetraacetic acid (EDTA) plasma samples were obtained for all LLS IPs and partners at inclusion. From these samples, metabolomics biomarker data was quantified using high-throughput nuclear magnetic resonance (NMR) spectroscopy provided by the Nightingale Health platform. Experimentation and application details of the Nightingale NMR platform has previously been described^52,53^. Moreover, the metabolic biomarkers measured using the nightingale platform were used in a variety of publications (overview can be found here: https://nightingalehealth.com/publications).

### Scanian Economic-Demographic Database

The Scanian Economic-Demographic Database (SEDD) is a longitudinal database covering five rural Scanian parishes and the city of Landskrona. It spans the period 1812-1967, with full coverage of the villages from 1812 and for Landskrona from 1904. The SEDD database was constructed using register-type data from catechetical examination registers and was updated with information on births, marriages, and deaths from church books. Unique person numbers were introduced in Sweden by 1947. Through these person numbers individuals can be followed in the national Swedish registration, introduced in 1967. Persons who out-migrated from the research region before the introduction of the person number were linked to the 1950 Census and the Swedish Death Index. The obtained person numbers were subsequently used to track individuals in the Swedish national register for the period 1968-2015. The link to the Swedish Death Index yielded ancestral death dates anywhere in Sweden even for individuals who out-migrated from the research region before the person number or nationwide register data were introduced. At present (2022), the SEDD database contains 920,159 unique individuals.

Index person (IP) identification for this study happened in subsequent steps (Supplementary table 5). First, from the entire SEDD data we identified all persons (from here: IPs) who were part of the national register data in the years 1990-1995 and between ages 45-60, and followed them in the national registers for the period 1990-2015. Second, IPs were selected to have known grandparents on at least one side of the family (maternal or paternal), and whose parents were from an extinct birth cohort (born before 1915) to ensure complete information about their date of death. Third, we included lifespan information of their parents, aunts and uncles, and their grandparents. Fourth, IPs who were found in the hospital records in the year preceding their eligibility for the study (1989-1994) were excluded to minimize the number of IPs with existing conditions receiving hospital treatments. Lastly, partners of IPs were excluded to ensure mutually exclusive ancestral information. In total, 1,493 Index Persons (IPs, F3), 2,969 parents (F2), 5,830 aunts and uncles (F2), and 3,028 grandparents (F1) were included in this study.

The Swedish hospital registers reached nationwide coverage in 1987 and records are considered complete from 1989. The main diagnosis for each hospitalization has been recorded in ICD-9 coding from 1987-1997 and ICD-10 coding 1997-2015. We recoded ICD-9 diagnoses to ICD-10 using the official crosswalk provided by Statistics Sweden. Diseases are specified identical to the LLS (Supplementary Table 4) to ensure comparability between the databases. It is relevant for our analyses to mention that only 214 IPs (8.6%) die without ever receiving a hospital diagnosis as a higher percentage would have warranted a competing risk analysis (see statistics section for more details).

### Lifetables

In the Netherlands and Sweden, population based cohort lifetables are available from 1850 and 1800 respectively, until 2021^54,55^. These lifetables contain, for each birth year and sex, an estimate of the hazard of dying between ages x and x + n (hx) based on yearly intervals (n=1) up to 99 years of age. Conditional cumulative hazards (Hx) and survival probabilities (Sx) can be derived using these hazards. In turn, we can determine to which sex and birth year based survival percentile each person of our study belonged to. For example: a person was born in 1876, was a female, and died at age 92. According to the lifetable information this person belonged to the top three percent survivors of her birth cohort, meaning that only three percent of the women born in 1876 reached a higher age. We used the lifetables to calculate the birth cohort and sex specific survival percentiles for all persons in the LLS and SEDD. This approach prevents against the effects of secular mortality trends over the last centuries and enables comparisons across study populations^56,57^. In SEDD, we focused only on extinct birth cohorts and death ancestors. However, In the LLS some ancestors (only aunts/uncles) were still alive (right censoring). To deal with non-extinct birth cohorts, we used the prognostic lifetables provided by Statistics Netherlands^54,55^ and to deal with right censoring we used single imputation where we estimated an age of death based on the remaining life expectancy at the age of censoring.

### Scores

The Longevity Relatives Count (LRC) score was used in LLS and SEDD to map the offspring’s family history of longevity. The LRC score indicates the proportion of ancestors that became long-lived, weighted by the genetic distance between IPs (and partners in LLS) and their ancestors. For example, an LRC of 0.5 indicates 50% long-lived ancestors. For this study, two generations of ancestors were available to calculate the LRC score for IPs and one generation for the partners of the LLS IPs (Figure 1 and Supplementary Figure 1). In the LLS, the LRC score was calculated using the mortality information as updated in 2021. In the SEDD, IPs were identified in such a way that all ancestors were deceased at the start of follow-up. The LRC score has been described in detail by van den Berg et al, 2020 in Aging Cell^23^. Supplementary Figure 6A-B depict the LRC score distribution in the LLS and SEDD. Additionally, in de LLS study, the MetaboHealth score was used as an indicator for the (metabolomic) health of the offspring and partners at study inclusion. This previously published score was generated by based on NMR metabolomics data in ∼40.000 European study participants and provides a weighted summary of 14 independent metabolites covering 5-10 years mortality risk and metabolite markers of lipid metabolism, fatty acid metabolism, glycolysis, fluid balance, and inflammation^26^. Supplementary Figure 6C depicts the MetaboHealth score distribution in the LLS.

### Statistical analyses

Statistical analyses were conducted using R version 4.0.2^58^. We reported 95% confidence intervals (CIs) and considered p-values statistically significant at the 5% level (α=0.05). A list of used R-packages and version numbers will be made available on gitlab (see code availability statement). In all random effect and frailty models we consider the F3 IPs who share the same parents as a family. Random effect and frailty models were used to adjust for within-family relations of the F3 IPs

#### Logistic and linear mixed model

To compare disease and medication prevalence between (LRC-based) LLS IPs and partners we fitted a logistic mixed model (1) and to compare the MetaboHealth score between the LRC groups (LRC_g1-3 with LRC_g4) in the LLS we fitted a linear mixed model (2):

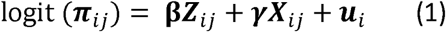

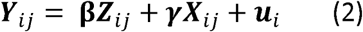

Where *Y*_*ij*_ is a vector of responses for person *j* in family *i*. and *π*_*ij*_ = *P*(*Y*_*ij*_ = 1|***Z***_*ij*_, ***X***_*ij*_, *u*_*i*_) when considering logistic regression. **β** is a vector of regression coefficients for the main effects of interest (***Z***). **γ** is a vector of regression coefficients for the effects of possible confounders (***X***). ***u*** is a vector of unobserved random effects shared by each member of the same family i and was assumed to follow a normal distribution. All analyses performed using logistic and linear mixed models have been adjusted for sex and age at study inclusion. In addition, the MetaboHealth score analyses have been adjusted for medication use.

#### Survival analysis (Cox-type random effects regression model)

To compare prospective disease incidences between (LRC-based) offspring and partners we fitted three different Cox-type random effects models:

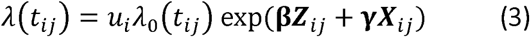

In the first type of models, we model t_ij_=age at first disease onset; in the second type of models the outcome of interest is t_ij_=age at the second disease onset (multi-morbidity onset). In both cases, the models are adjusted for left truncation given by the age at entry in the study. In the third type of models, we consider the time between the first and the second disease onset; i.e. t_ij_=age at second disease onset where age at the first disease onset is considered as the left-truncation time in this analysis. *λ*_0_(*t*_*ij*_) refers to the baseline hazard, which is left unspecified in a Cox-type model. **β** is the vector of regression coefficients for the main effects of interest (***Z***). **γ** is a vector of regression coefficients for the effects of possible confounders (***X***). *u*_*i*_ > 0 refers to an unobserved random effect (frailty) shared by the members of the same family i and was assumed to follow a gamma distribution. All survival models were adjusted for sex. Additionally, the third type of analysis focusing on the time from first to second disease (Table 3) has been further adjusted for age at study inclusion as we did not limit our sample to persons without any diseases at the start of follow-up.

## Ethical regulations

Leiden Longevity Study: In accordance with the Declaration of Helsinki, we obtained informed consent from all participants prior to their entering the study. Good clinical practice guidelines were maintained. The study protocol was approved by the ethical committee of the Leiden University Medical Center before the start of the study (P01.113).

SEDD: The SEDD has approval for research from Regionala etikprövningsnämnden, Lund, (dnr 161/2006, dnr 627/2010), and instructions from Datainspektionen, Stockholm (dnr 1999-2005).

## Author contributions

Niels van den Berg is the study investigator and was responsible for initiating the study, data management, data analyses, writing the manuscript and finalizing it. Ingrid van Dijk was responsible for the data organization and analyses of the Swedish data. Mar Rodriguez-Girondo provided overall support on statistical analyses. P. Eline Slagboom and Marian Beekman provided overall coordination and supervision.

## Code availability

The scripts containing the code for data pre-processing and data analyses can be freely downloaded at: https://git.lumc.nl/publications/longevity-family-diseases

## Data availability

The individual-level data from the SEDD, the Statistics Sweden, and LLS are protected by Swedish and Dutch personal integrity laws, and other (privacy) regulations. As such, restrictions apply to the availability of the LLS and SEDD data, which were used under license for the current study, and so are not publicly available. For both datasets, summary statistics are available upon request to the corresponding author (Niels van den Berg). The LLS data is available for replication purposes upon reasonable request to P. Eline Slagboom and if replication is conducted within the secure LUMC network environment. Researchers can gain access to the SEDD data as used in this study if relevant permissions have been obtained in accordance with the restrictions stated by the Regional Ethical Review Board, the Swedish Data Inspection Board and Lund University.

## Acknowledgments

The construction and maintenance of the LLS data has received funding from the European Union’s Seventh Framework Programme (FP7/2007-2011) under grant agreement number 259679. This study was further supported by the Innovation-Oriented Research Program on Genomics (SenterNovem IGE05007), the Centre for Medical Systems Biology and the Netherlands Consortium for Healthy Ageing (grant 050-060-810), all in the framework of the Netherlands Genomics Initiative, Netherlands Organization for Scientific Research (NWO), by BBMRI-NL, a Research Infrastructure financed by the Dutch government (NWO 184.021.007 and 184.033.111). Ingrid van Dijk was funded through the research program “Landskrona Population Study” and Niels van den Berg and Ingrid van Dijk were funded through the research project “An Age Old Advantage?” (P21-0139), the Swedish Foundation for the Humanities and Social Sciences (Riksbankens Jubileumfond, RJ). Niels van den Berg was further funded by the Netherlands Organization for Scientific Research, domain Health Research and Medical Sciences (09120012010052). We thank Bianca Schutte, Rinske van Reijen, and Yotam Raz for digitizing and organizing the LLS medication and disease data.

