## Supplementary for "Increasing number of long-lived ancestors associates with up to a decade of healthspan extension and a healthy metabolomic profile in mid-life"

**Supplementary Table 1: Time from study inclusion to first disease and multimorbidity**

|  | **A: Time from inclusion to first disease** | | | | **B: Time from inclusion to 2 diseases** | | | |
| --- | --- | --- | --- | --- | --- | --- | --- | --- |
|  | **N (prop)** | **Events**  **(prop)** | **HR (95% CI)** | **P-Value** | **N (prop)** | **Events**  **(prop)** | **HR (95% CI)** | **P-Value** |
| **Age related diseases** |  |  |  |  |  |  |  |  |
| Groups |  |  |  |  |  |  |  |  |
| Offspring | 917 (0.70) | 362 (0.39) | 0.79 (0.65-0.97) | 2.32·10^-2^ | 611 (0.70) | 73 (0.12) | 0.55 (0.36-0.85) | 6.83·10^-3^ |
| Partners (ref) | 395 (0.30) | 171 (0.43) |  |  | 268 (0.30) | 47 (0.18) |  |  |
| Sex |  |  |  |  |  |  |  |  |
| Females | 733 (0.56) |  | 0.79 (0.66-0.94) | 8.78·10^-3^ | 611 (0.70) |  | 0.59 (0.40-0.89) | 1.04·10^-2^ |
| males (ref) | 579 (0.44) |  |  |  | 268 (0.30) |  |  |  |
| **Metabolic diseases** |  |  |  |  |  |  |  |  |
| Groups |  |  |  |  |  |  |  |  |
| Offspring | 917 (0.70) | 261 (0.23) | 0.71 (0.55-0.90) | 5.18·10^-3^ | 611 (0.70) | 45 (0.07) | 0.51 (0.26-0.97) | 3.96·10^-2^ |
| Partners (ref) | 395 (0.30) | 135 (0.34) |  |  | 268 (0.30) | 29 (0.11) |  |  |
| Sex |  |  |  |  |  |  |  |  |
| Females | 733 (0.56) |  | 0.84 (0.68-1.05) | 1.26·10^-1^ | 611 (0.70) |  | 0.52 (0.30-0.92) | 2.41·10^-2^ |
| males (ref) | 579 (0.44) |  |  |  | 268 (0.30) |  |  |  |
| **Malignancies** |  |  |  |  |  |  |  |  |
| Groups |  |  |  |  |  |  |  |  |
| Offspring | 917 (0.70) | 130 (0.14) | 0.95 (0.70-1.31) | 7.66·10^-1^ | 611 (0.70) | 7 (0.01) | 1.39 (0.29-6.70) | 6.82·10^-1^ |
| Partners (ref) | 395 (0.30) | 56 (0.14) |  |  | 268 (0.30) | 2 (0.01) |  |  |
| Sex |  |  |  |  |  |  |  |  |
| Females | 733 (0.56) |  | 0.66 (0.49-0.88) | 4.70·10^-3^ | 611 (0.70) |  | 1.06 (0.28-3.96) | 9.30·10^-1^ |
| Males (ref) | 579 (0.44) |  |  |  | 268 (0.30) |  |  |  |

Table shows the time from inclusion to first disease in panel A, and the time from inclusion to having 2 diseases (panel B). N is the group size used for the analyses and prop. Is the proportion from the total. Events are the events of the specific diseases, for example age related diseases, and prop. indicates the proportion from the size of a specific group (LLS IPs or partners). HR is the abbreviation for Hazard Ratio. Statistical testing was performed using Wald tests for the conditional log-hazard ratio estimated with a Cox-type frailty regression model. The analyses are adjusted for sex, different ages of study entry (left truncation) and right censoring. Survival curve details can be found in Supplementary Figure 3 and 4. To study time from inclusion to first disease, only persons without any disease at inclusion were studied. Note that the numbers in panel B are lower than those in panel A. This is because the censoring group reflects those for whom we have not observed any disease at the end of follow-up. As a results, persons with only one disease are excluded from the analyses.

**Supplementary Table 2: Time from first disease to multimorbidity**

|  | **Time from first disease to second disease** | | | |
| --- | --- | --- | --- | --- |
|  | **N (mean)** | **Events (mean)** | **HR (95% CI)** | **P-Value** |
| **Age related 🡪 age related diseases** |  |  |  |  |
| Groups |  |  |  |  |
| Offspring | 500 (0.68) | 79 (0.16) | 0.46 (0.26-0.83) | 9.82·10^-3^ |
| Partners (ref) | 237 (0.32) | 55 (0.23) |  |  |
| Sex |  |  |  |  |
| Females | 345 (0.47) |  | 0.45 (0.27-0.73) | 1.19·10^-3^ |
| males (ref) | 392 (0.53) |  |  |  |
| Age at inclusion | 737 (60) |  | 0.92 (0.86-0.96) | 3.49·10^-3^ |
| **Age related 🡪 Metabolic diseases** |  |  |  |  |
| Groups |  |  |  |  |
| Offspring | 500 (0.68) | 62 (0.12) | 0.33 (0.14-0.81) | 1.46·10^-2^ |
| Partners (ref) | 237 (0.32) | 44 (0.19) |  |  |
| Sex |  |  |  |  |
| Females | 345 (0.47) |  | 0.41 (0.21-0.79) | 7.47·10^-3^ |
| males (ref) | 392 (0.53) |  |  |  |
| Age at inclusion | 737 (60) |  | 0.89 (0.82-0.96) | 3.92·10^-3^ |
| **Age related 🡪 Malignancies** |  |  |  |  |
| Groups |  |  |  |  |
| Offspring | 500 (0.68) | 17 (0.03) | 0.58 (0.27-1.25) | 1.66·10^-1^ |
| Partners (ref) | 237 (0.32) | 11 (0.05) |  |  |
| Sex |  |  |  |  |
| Females | 345 (0.47) |  | 0.58 (0.26-1.28) | 1.74·10^-1^ |
| Males (ref) | 392 (0.53) |  |  |  |
| Age at inclusion | 737 (60) |  | 0.89 (0.79-0.99) | 4.02·10^-2^ |

Table shows the time from first to a specific second disease. N is the group size and prop. Is the proportion from the total. Events are the events of the specific disease, for example age related diseases, and prop. indicates the proportion from size of a specific group (offspring or partners). Sd indicates the standard deviation. HR is the abbreviation for Hazard Ratio. Statistical testing was performed using Wald tests for the conditional log-hazard ratio estimated with a frailty Cox-type frailty regression model. The analyses are adjusted for sex and age at inclusion.

**Supplementary Table 3: MetaboHealth score differences for LRC groups at LLS study inclusion**

|  | **N (mean)** | **Beta estimate (95% CI)** | **P- Value** |
| --- | --- | --- | --- |
| Groups |  |  |  |
| Offspring LRC10_60% | 121 (0.06) | -0.098 (-0.184 / -0.012) | 2.66*10^-2 |
| Original participants | 1297 (0.67) | -0.032 (-0.077 / 0.012) | 1.55*10^-1 |
| Partners with ≥ top 10% parent | 135 (0.07) | -0.016 (-0.091 / 0.058) | 6.61·10^-1 |
| Partners without top 10% parents (ref) | 397 (0.20) |  |  |
| Sex |  |  |  |
| Females | 1083 (0.56) | 0.004 (-0.039 / 0.029) | 7.76*10^-1 |
| males (ref) | 867 (0.44) |  |  |
| Age at inclusion | 1950 (59.20) | 0.007 (0.005 / 0.010) | 1.37*10^-7 |
| Medication use |  |  |  |
| Yes | 980 (0.50) | 0.130 (0.096 / 0.164) | 1.36e-13 |
| No (ref) | 970 (0.50) |  |  |

In this table the metabolome of the LRC groups are compared at the moment of study inclusion using the MetaboHealth score. N indicates the total group size. Mean indicates a proportion for categorical measurements and the group average for continuous measurements. Estimate indicates the beta coefficient of the linear regression analysis. Statistical testing was performed using T-tests for the Beta coefficients estimated with a mixed-model linear regression analysis. The analyses are adjusted for sex, age at inclusion, and medication use.

**Supplementary table 4: Disease groups based on ICD scores for GP data**

| **Diseases (ICD10-code)** | **Age related** | **Metabolic** | **Malignancies** |
| --- | --- | --- | --- |
| TIA (I60-I69) | X | X |  |
| CVA (I63) | X | X |  |
| AP (I20) | X | X |  |
| MI (I21) | X | X |  |
| Hypertension (I10) | X | X |  |
| Diabetes (E10–E14) | X | X |  |
| Prostate (C61) | X |  | X |
| Mamma (C50) | X |  | X |
| Lung (C34) | X |  | X |
| Colon (C18-C20) | X |  | X |
| Other (C00-D48) | X |  | X |

Table provides an overview of the diseases, their ICD10 codes, and the disease clusters.

**Supplementary Table 5: Inclusion and exclusion criteria of the SEDD sample group**

|  | **Number of unique remaining IDs** | **Number of unique IDs removed** |
| --- | --- | --- |
| All persons in SEDD and national registers | 920,195 |  |
| Parents are known (parent ids are known) | 369,313 | -550,882 |
| At least one set of grandparents known (maternal or paternal ids are known) | 180,769 | -188,544 |
| Observed in the years 1990-1995 | 121,967 | -58,802 |
| Age 45-60 in the years 1990-1995 | 4,831 | -117,136 |
| Parents from extinct cohort (born < 1915) | 2,732 | -2,099 |
| Removal of individuals who are inpatients in 1989 | 2,500 | -232 |
| Other data cleaning |  | -3 |
| Total number of Index Persons | 2,497 |  |

Table provides an overview of the SEDD inclusion and exclusion criteria.

**Supplementary Figure 1: LLS study design**

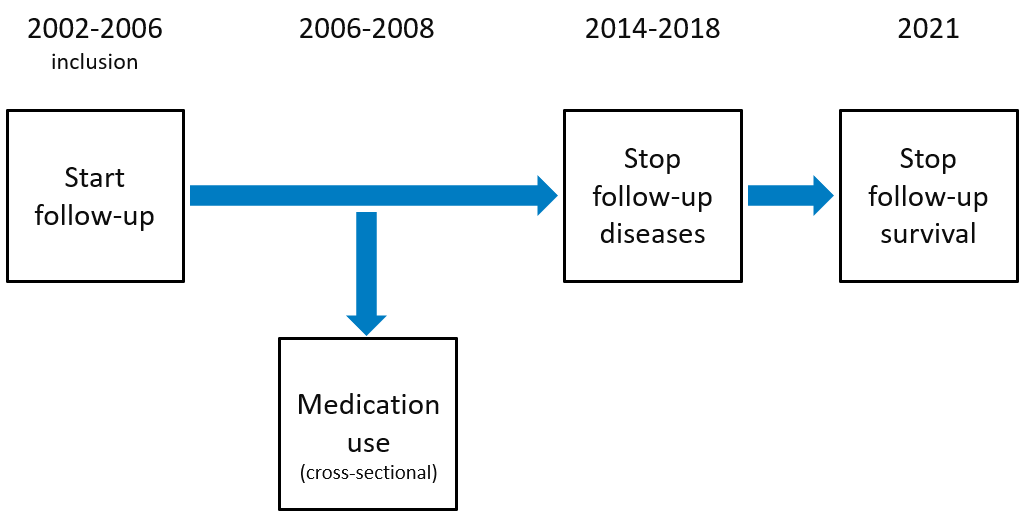

This figure illustrates the Leiden Longevity Study (LLS) design. The LLS was initiated between 2002-2006. Subsequently data on medication use was available for the period 2006-2008. Disease incidence data was available from follow-up until 2018 and mortality data was available from follow-up until 2021.

**Supplementary Figure 2: Survival curves corresponding to Table 2A**

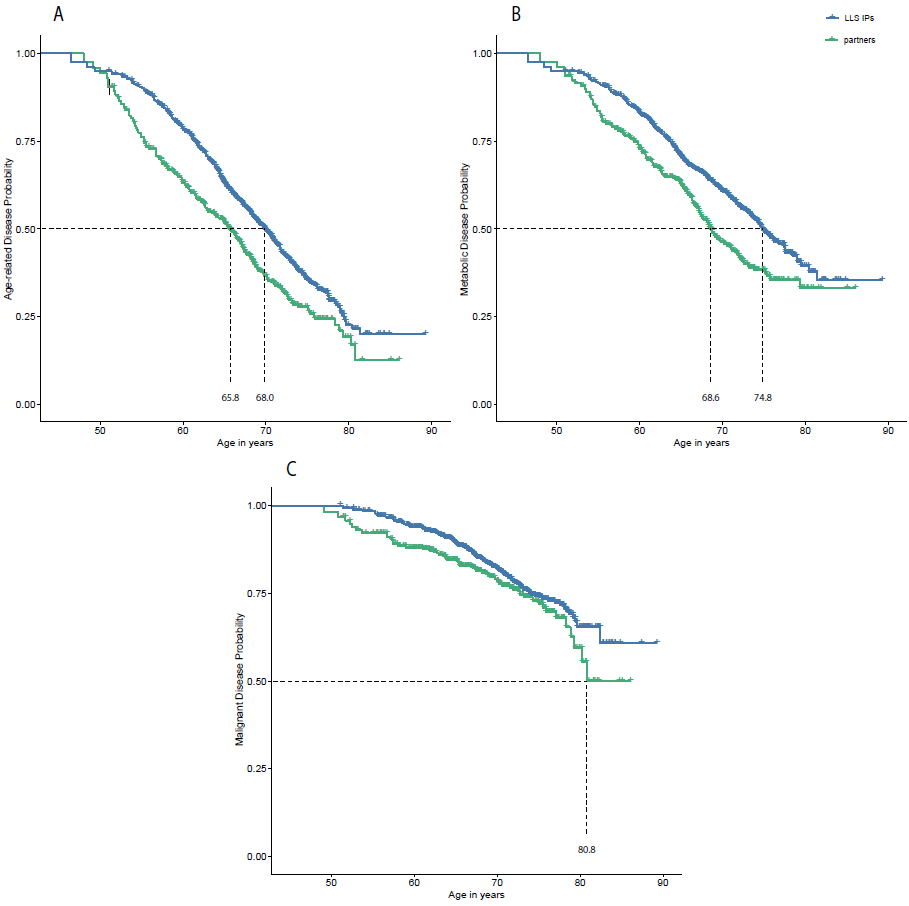

This figure depicts survival curves reflecting age-related (Panel A), metabolic (Panel B), and malignant (Panel C) disease incidence within the Leiden Longevity Study (LLS). The x-axis show age in years and the y-axis show disease incidence. Dotted lines represent the age at which 50% of the members of a specific group had their first disease. The blue line represents LLS Index Persons (IPs) and the green line represents their partners. Vertical lines within the colored lines represent right censoring events. Survival curves are adjusted for left truncation and right censoring.

**Supplementary Figure 3: Survival curves corresponding to Table 2B**

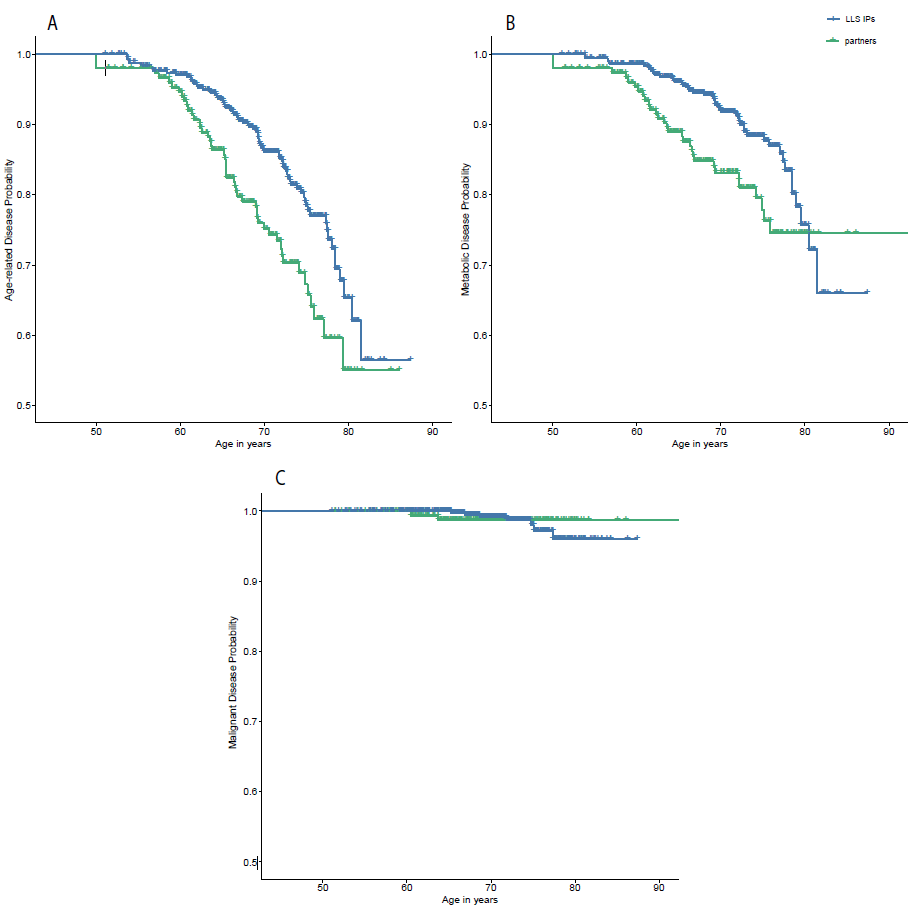

This figure depicts survival curves reflecting age-related (Panel A), metabolic (Panel B), and malignant (Panel C) disease incidence within the Leiden Longevity Study (LLS). The x-axis show age in years and the y-axis show disease incidence. Dotted lines represent the age at which 50% of the members of a specific group had 2 diseases. The blue line represents LLS Index Persons (IPs) and the green line represents their partners. Vertical lines within the colored lines represent right censoring events. Survival curves are adjusted for left truncation and right censoring.

**Supplementary Figure 4: Survival curves corresponding to Table 2C**

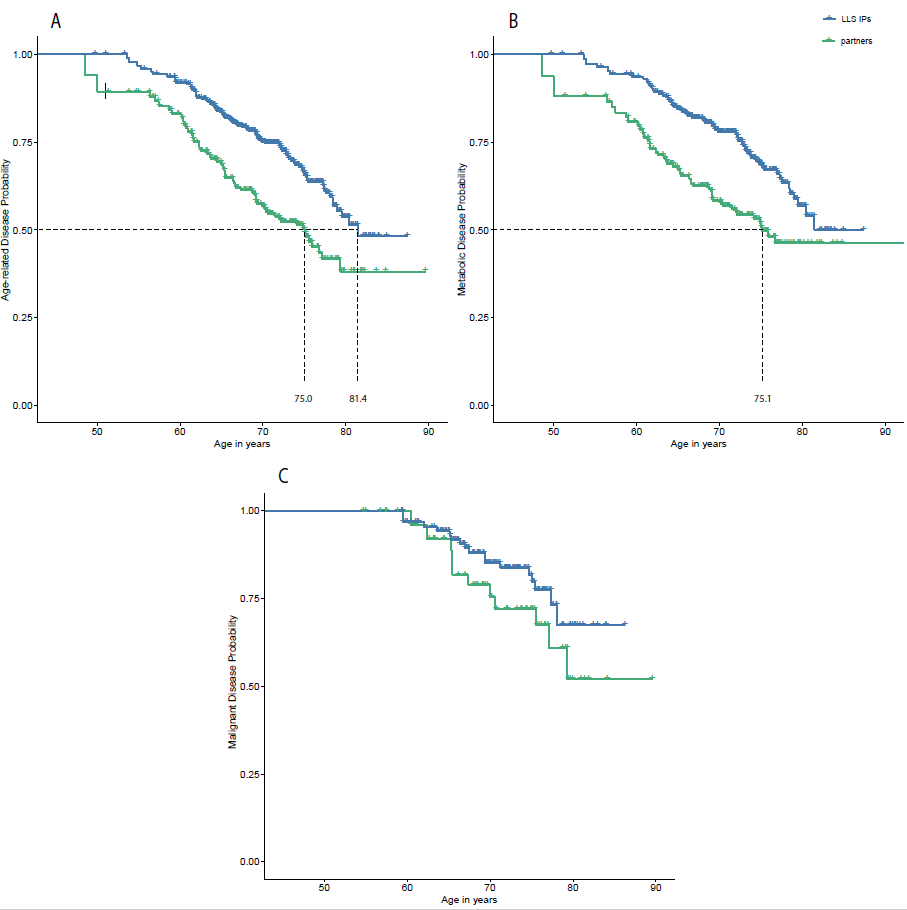

This figure depicts survival curves reflecting age-related (Panel A), metabolic (Panel B), and malignant (Panel C) disease incidence within the Leiden Longevity Study (LLS). The x-axis show age in years and the y-axis show disease incidence. Dotted lines represent the age at which 50% of the members of a specific group had their second disease after already being diagnosed with a first disease. The blue line represents LLS Index Persons (IPs) and the green line represents their partners. Vertical lines within the colored lines represent right censoring events. Survival curves are adjusted for left truncation and right censoring.

**Supplementary Figure 5: Survival curves corresponding to Table 3**

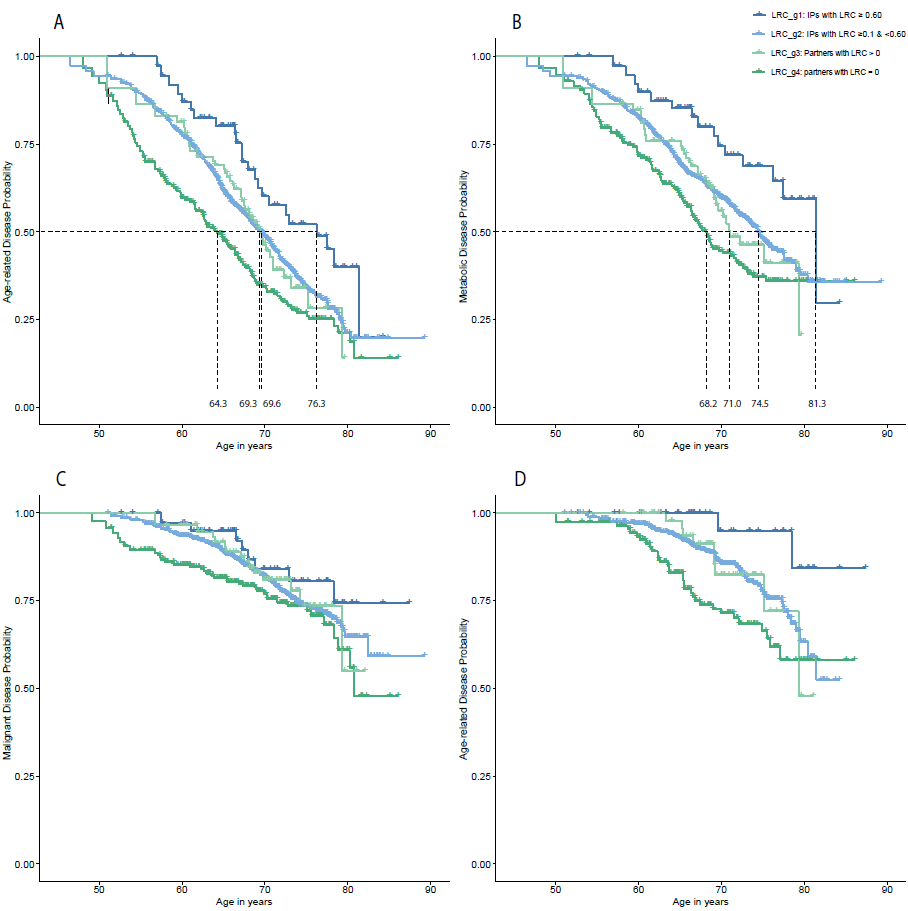

This figure depicts survival curves reflecting age-related (Panel A), metabolic (Panel B), malignant (Panel C), and age-related (Panel D) disease incidence within the Leiden Longevity Study (LLS). The x-axis show age in years and the y-axis show disease incidence. Dotted lines represent the age at which 50% of the members of a specific group had their first disease (Panel A-C) and the age at which 50% of the members of a specific group had 2 diseases. The blue line represents LLS Index Persons (IPs) and the green line represents their partners. Vertical lines within the colored lines represent right censoring events. Survival curves are adjusted for left truncation and right censoring.

**Supplementary Figure 6: Distribution of LRC and MetaboHealth scores**

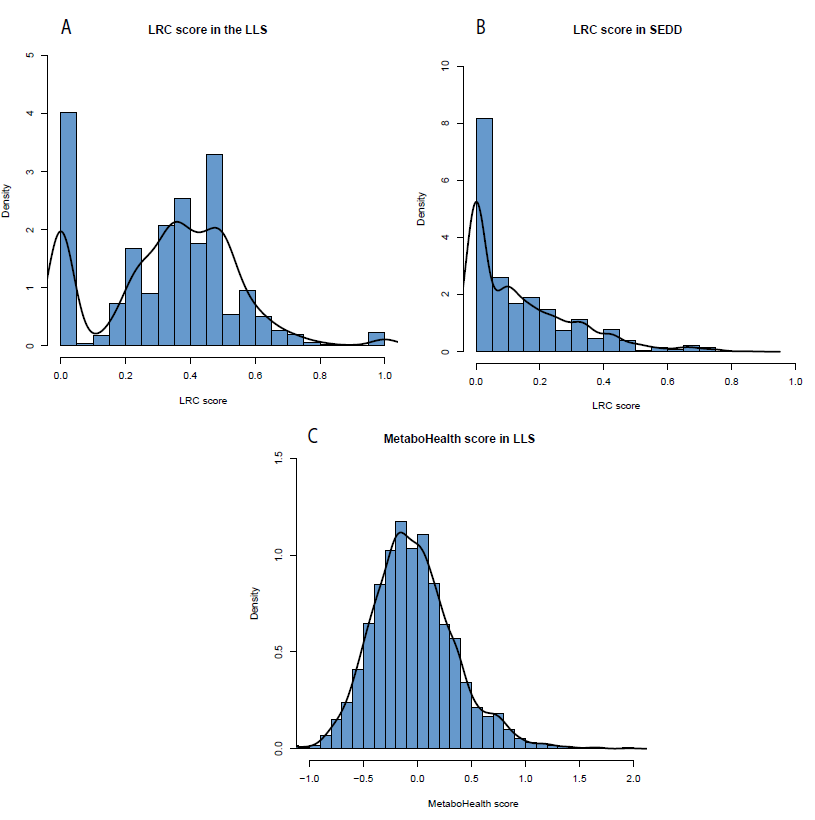

Figure depicts the distribution of the LRC score in the LLS (Panel A) and SEDD (Panel B). It further depicts the distribution of the MetaboHealth score in the LLS (Panel C)
